# Single day construction of multi-gene circuits with 3G assembly

**DOI:** 10.1101/260851

**Authors:** Andrew D. Halleran, Anandh Swaminathan, Richard M. Murray

## Abstract

The ability to rapidly design, build, and test prototypes is of key importance to every engineering discipline. DNA assembly often serves as a rate limiting step of the prototyping cycle for synthetic biology. Recently developed DNA assembly methods such as isothermal assembly and type IIS restriction enzyme systems take different approaches to accelerate DNA construction. We introduce a hybrid method, Golden Gate-Gibson (3G), that takes advantage of modular part libraries introduced by type IIS restriction enzyme systems and isothermal assembly‘s ability to build large DNA constructs in single pot reactions. Our method is highly efficient and rapid, facilitating construction of entire multi-gene circuits in a single day. Additionally, 3G allows generation of variant libraries enabling efficient screening of different possible circuit constructions. We characterize the efficiency and accuracy of 3G assembly for various construct sizes, and demonstrate 3G by characterizing variants of an inducible cell-lysis circuit.

## Introduction

The ability to rapidly design, build, and test genetic circuits is of crucial importance to synthetic biology. Despite advances in rapid *in vitro* prototyping popularized by cell-free systems, *in vivo* testing often requires laborious construction of large genetic constructs onto a small number of plasmids.^1,2^ This assembly process often serves as a rate limiting step in iterative design, and multiple DNA assembly methods have been developed to address the bottleneck.^3-11^

Currently, two of the most popular assembly methods are Gibson assembly and systems based on type IIS restriction enzymes.^4-8^ While both methods can be used to successfully construct complex DNA constructs, they also have significant drawbacks. Cloning strategies based on type IIS restriction systems rely on hierarchical cloning schemes that introduce intermediate constructs that must be cloned and amplified before assembling the final desired sequence.^4-8^ Despite this, the modular part-level design and inherent nature to readily build combinatorial circuits has made it a popular choice in the synthetic biology community.

Gibson assembly requires design and synthesis of oligonucleotide sequences that are unique to the target and have long overhangs to adjacent fragments.^9^ Not only does this increase the cost for generating each construct, it also introduces a delay in the design-build-test cycle. However, Gibson assembly does have several benefits including the ability to combine many fragments in a single reaction and resulting constructs that are scarless, lacking unnecessary sequences frequently left behind by restriction enzyme-based systems.

We develop a hybrid Golden Gate-Gibson (3G) assembly method that leverages the advantages of both type IIS and isothermal assembly to enable construction of multi-gene circuits in a single day. This method builds upon the existing CIDAR MoClo standardized part library and unique nucleotide sequences used with Gibson assembly to produce high-efficiency assembly of entire genetic circuits a single day.^7^

A similar approach has recently been proposed by Pollak *et al*.^8^ Their approach deposits Golden Gate products into intermediate vectors that can then be amplified via PCR and used in a subsequent Gibson reaction. This approach requires construction of all possible intermediate vectors, and results in a multi-day assembly pipeline. Our method bypasses the need for intermediate vectors and can be performed in a single day. In addition, we show here that random libraries can be created and characterize both the efficiency and accuracy of 3G assembly. We demonstrate 3G by building and testing variants of an inducible cell lysis circuit.

## Results

Golden Gate assembly allows for the combination of standardized part-level components (e.g. promoters, UTRs, CDSs, and terminators) into a single transcriptional unit.^4-8^ By using a common type IIS restriction enzyme to generate standardized overhangs, libraries of parts can be built and re-used across different designs. However, these approaches typically rely on a hierarchical assembly model in which part-level components are first deposited into “level I” part vectors, and then combined into individual transcriptional units (TUs) in a “level II” vector. Level II vectors, each containing an individual TU, can then be combined into a “level III” vector, bringing together multiple independent transcriptional units into a single plasmid backbone. The intentional hierarchy of this system is flexible, but requires an intermediate step in the transition from part-level components to entire circuits, thus increasing the time to build a desired construct.

A popular alternative method for DNA construction is isothermal assembly.^9^ Isothermal assembly requires long regions (≥ 15bp) of homology between adjacent fragments to join them together. While this approach allows for the construction of large stretches of DNA with high efficiency and accuracy, addition of regions of homology via overhang PCR requires custom design and synthesis of oligonucleotides for each assembly.

3G assembly takes advantage of both the standardized part libraries provided by Golden Gate and the ability of isothermal assembly to efficiently stitch together large constructs (Figure 1a). To bridge the two assembly strategies, we developed adapter sequences based on unique nucleotide sequence (UNS) that ligate to and flank each transcriptional unit (1b).^10,11^ By adding the standard BsaI recognition site and overhangs, part-level Golden Gate reactions also incorporate a common landing site for PCR and isothermal assembly. Following the Golden Gate reactions, the assembled product is used as template for a PCR reaction that primes off the adapters added to each side of the transcriptional unit. These fragments can then be combined in a standard Gibson reaction with a backbone containing terminal acceptor UNS sequences (UNS1 and UNS10) at either end. This pipeline takes less than 8 hours with minimal hands-on time and generates entire circuits (Figure 1a-b).

**Figure 1:**
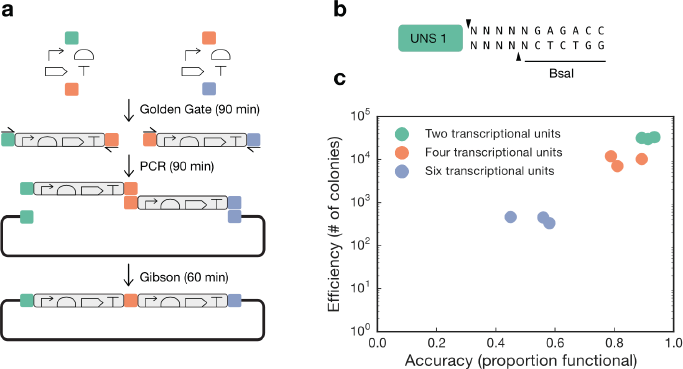
3G assembly overview. (A) Pipeline for 3G assembly. A Golden Gate reaction is performed to assemble each transcriptional unit of interest flanked by UNS sequences (colored boxes). The Golden Gate product is then amplified with primers specific to each UNS sequence. These PCR products are then combined with a backbone containing flanking UNS (1 and 10) sequences and assembled using Gibson assembly. (B) Schematic of the UNS adapter. The entire 40bp unique nucleotide sequence is followed by a programmable sticky-end generated by BsaI. This design allows extension of the UNS adapter method to other existing Golden Gate systems. (C) Assembly and accuracy for 3G assemblies using two, four, or six transcriptional units. Efficiency is the number of colonies generated by a single transformation of 50μL of competent cells. Accuracy is the fraction of colonies that were functionally validated. Each dot represents an independent Golden Gate, PCR, Gibson, and transformation and represents 48 colonies screened for accuracy.

3G assembly not only provides a single day pipeline for constructing entire circuits, it is also highly efficient and accurate. To quantify the accuracy and efficiency of assemblies with different numbers of transcriptional units we designed assemblies with two, four, or six transcriptional units (Figure 1c). To enable functional screening, each set of two transcriptional units consist of a constitutively expressed transcription factor (TF) and a distinct fluorescent protein driven by that transcription factor. Assemblies with two inserts consist of pSal-BCD2-sfCFP- ECK120033736 and J23106-BCD12-NahR- ECK120033736, and should be activated by addition of salicylate.^12^ Assemblies with four inserts contain the NahR / pSal system in addition to pLas-BCD2-sfYFP-ECK120033736 and J23106-BCD12-LasR-ECK120033736 and should induce both sfCFP and sfYFP expression upon addition of salicylate and 3OC12 AHL. Finally, six insert assemblies contain the NahR / pSal system, LasR / pLas system, along with pTet-BCD2-mScarletI-ECK120033736 and J23106-BCD12-TetR-ECK120033736 and should induce sfCFP, sfYFP, and mScarlet expression upon addition of salicylate, 3OC12 AHL, and anhydrotetracycline. Two insert assemblies have > 90% accuracy and six insert assemblies have > 50% accuracy (Figure 1c). Standard transformation into commercially available competent cells generates several hundred (six insert assembly) to tens of thousands (two insert assembly).

The high efficiency and accuracy of 3G enables construction and screening of circuit variants. By including pools of multiple different part-level components with the same overhangs in the Golden Gate reaction, the final circuit plasmid will contain many possible variations of part combinations. Therefore, 3G can be used to create large libraries of entire circuits that vary in part-level components. This enables rapid generation and screening of circuits variants.

To demonstrate the effectiveness of the 3G assembly method, we built variants of an inducible cell lysis circuit in which induction of the phage toxin ΦX174E drives cell lysis (Figure 2a).^13^ To tune the properties of the response profile we varied the 5’ UTR controlling ΦX174E expression, and the promoter and 5’ UTR that constitutively express the transcription factor. To screen circuit behavior we selected 32 colonies at random. Colonies were grown for 4 hours in M9CA + chloramphenicol, then diluted into fresh media either containing 0μM Las AHL or 1μM Las AHL (Figure 2b).^14^ Not only did we identify both responsive and non-responsive colonies, but we also identified a critical design parameter. Colonies that showed a low maximum OD600 in the uninduced condition uniformly had strong 5’ UTRs driving ΦX174E, leading us to suspect leak in the ΦX174E gene caused spontaneous lysis, even in the absence of inducer molecule. This insight emphasizes the importance of screening multiple different possible designs. To discern dynamics at the single cell level we performed time-lapse microscopy on a responsive colony (Figure 2c). We performed Sanger sequencing and identified that we constructed and tested 20 unique variants (Figure 2d).

**Figure 2:**
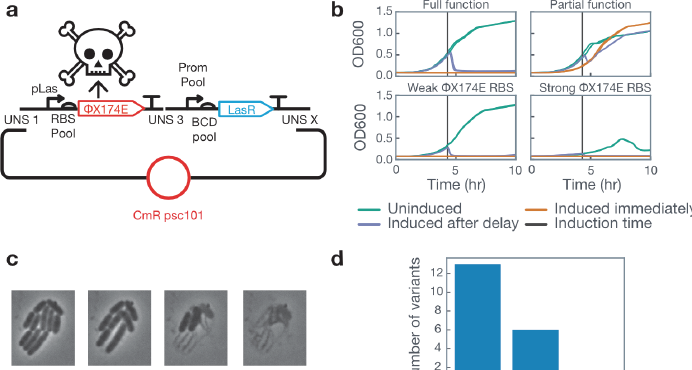
Inducible cell-lysis circuit variant characterization. (A) Circuit diagram. LasR is constitutively expressed and can be induced to activate expression of the ΦX174E lysis protein. To screen different promoter and 5’ UTR combinations we used a pool of different constitutive promoters and BCDs. The promoter pool used for constitutive LasR, BCD pool used for LasR, and the RBS pool used for ΦX174E can all be found in Supplementary Table 2. (B) 32 colonies were screened for function by growing colonies and inducing immediately, inducing after ~4 hours of growth, or not inducing. Fully functional colonies displayed correct behavior (no growth in immediate induction, immediate drop in OD in induced after delay, robust growth in uninduced), partial function colonies did not respond as desired to induction of the ΦX174E protein. Colonies that showed a lower OD in the uninduced condition uniformly had a strong RBS driving expression of ΦX174E. All traces show two lines, each corresponding to a replicate culture. (C) Time-lapse microscopy of a responsive colony displayed cell-lysis. (D) Sanger sequencing revealed that we screened 20 unique circuit variants.

By combining single-day assembly with the ability to generate libraries of circuit variants, 3G assembly addresses two critical aspects of the design-build-test cycle for synthetic biology. We believe that 3G will vastly reduce the time required to go from a design on the board to a functional circuit in a cell.

## Materials and methods

### 3G assembly

A detailed assembly protocol can be found in the Supplementary Information. All primers and adapter sequences are available in Supplementary Table S1. Golden Gate reactions were performed as described by Iverson *et al*.^7^ All parts were used at a final concentration of 3nM; annealed adapters were used at a final concentration of 5nM. To amplify the Golden Gate product, 1.5μL of the reaction was used as template in a 50μL PCR reaction (NEB Q5 Hot Start High-Fidelity 2X Master Mix). PCR products were purified using the QIAquick Gel Extraction kit. Isothermal assembly was performed using NEBuilder HiFi DNA Assembly Master Mix. All fragments were combined at 2nM final concentration and incubated at 50°C for 1 hour. 2μL of the isothermal assembly reaction was then transformed into 50μL of chemically competent DH5α cells and grown overnight at 37°C (NEB 5-alpha Competent *E. coli* (High Efficiency)).

### Functional validation of 3G constructs

Colonies were picked into M9CA (Teknova) and grown at 37°C for 2 hours. Cultures were then diluted 1:100 into fresh M9CA + chloramphenicol (34μg/mL) or M9CA + chloramphenicol (34μg/mL) containing 100μM salicylate, 1μM 3OC12 AHL, and 100ng/mL anhydrotetracycline. Cells were then grown for eight hours at 37°C and final OD600 and fluorescence measurements were taken on a BioTek Synergy H1m. sfCFP was measured at 440nm / 480nm. sfYFP was measured at 502nm / 530nm. mScarlet was measured at 570nm / 600nm.

### Cell-lysis circuit characterization

Cell-lysis circuit characterization was performed in the JS006 *E. coli* strain. Colonies were picked into M9CA (Teknova) + chloramphenicol (34μg/mL) and grown at 37°C for 2 hours. Cultures were then split into three conditions each with two replicates. A no induction (0uM AHL) control, an immediate induction (1μM Las AHL) and a delayed induction (3-hour outgrowth followed by addition of 1μM Las AHL. All cultures were diluted 1:50 into M9CA + chloramphenicol (34μg/mL). Cultures were grown in a BioTek Synergy H1m at 37°C with OD600 readings taken every 5 minutes. Movies were performed as previously described.^15^ Briefly, uninduced cells in log phase were plated on agar pads and induced with 1μM Las AHL at T=0 minutes. Cells were grown in an incubated chamber at 37°C and bright field images were taken every 2 minutes on an Olympus IX81 inverted microscope.

## Acknowledgments

pSal, pLas, NahR, and LasR plasmids were generously provided by Adam Meyer. Plasmid vectors were provided by Douglas Densmore at the Cross-disciplinary Integration of Design Automation Research lab (Addgene Kit # 1000000059). The project depicted was sponsored by the Defense Advanced Research Projects Agency (Agreement HR0011-17-2-0008). The content of the information does not necessarily reflect the position or the policy of the Government, and no official endorsement should be inferred.

## References

1. Shin, J., and Noireaux, V. (2012) An E. coli cell-free expression toolbox: application to synthetic gene circuits and artificial cells, ACS Synthetic Biology 1, 29–41.

2. Sun, Z. Z., Yeung, E., Hayes, C. A., Noireaux, V., and Murray, R. M. (2013) Linear DNA for rapid prototyping of synthetic biological circuits in an Escherichia coli based TX-TL cell-free system, ACS Synthetic Biology 3, 387–397.

3. Shetty, R. P., Endy, D., and Knight, T. F. (2008) Engineering BioBrick vectors from BioBrick parts, Journal of Biological Engineering 2, 5.

4. Engler, C., Kandzia, R., and Marillonnet, S. (2008) A one pot, one step, precision cloning method with high throughput capability, PloS One 3, e3647.

5. Engler, C., Gruetzner, R., Kandzia, R., and Marillonnet, S. (2009) Golden gate shuffling: a one-pot DNA shuffling method based on type IIs restriction enzymes, PloS One 4, e5553.

6. Sarrion-Perdigones, A., Falconi, E. E., Zandalinas, S. I., Juárez, P., Fernández-del-Carmen, A., Granell, A., and Orzaez, D. (2011) GoldenBraid: an iterative cloning system for standardized assembly of reusable genetic modules, PloS One 6, e21622.

7. Iverson, S. V., Haddock, T. L., Beal, J., and Densmore, D. M. (2015) CIDAR MoClo: improved MoClo assembly standard and new E. coli part library enable rapid combinatorial design for synthetic and traditional biology, ACS Synthetic Biology 5, 99–103.

8. Pollak, B., Cerda, A., Delmans, M., Álamos, S., Moyano, T., West, A., Gutiérrez, R. A., Patron, N., Federici, F., and Haseloff, J. (2018) Loop Assembly: a simple and open system for recursive fabrication of DNA circuits, bioRxiv, 247593.

9. Gibson, D. G., Young, L., Chuang, R.-Y., Venter, J. C., Hutchison III, C. A., and Smith, H. O. (2009) Enzymatic assembly of DNA molecules up to several hundred kilobases, Nature Methods 6, 343.

10. Torella, J. P., Boehm, C. R., Lienert, F., Chen, J.-H., Way, J. C., and Silver, P. A. (2013) Rapid construction of insulated genetic circuits via synthetic sequence-guided isothermal assembly, Nucleic Acids Research 42, 681–689.

11. Torella, J. P., Lienert, F., Boehm, C. R., Chen, J.-H., Way, J. C., and Silver, P. A. (2014) Unique nucleotide sequence–guided assembly of repetitive DNA parts for synthetic biology applications, Nature Protocols 9, 2075.

12. Chen, Y.-J., Liu, P., Nielsen, A. A., Brophy, J. A., Clancy, K., Peterson, T., and Voigt, C. A. (2013) Characterization of 582 natural and synthetic terminators and quantification of their design constraints, Nature Methods 10, 659.

13. Scott, S. R., Din, M. O., Bittihn, P., Xiong, L., Tsimring, L. S., and Hasty, J. (2017) A stabilized microbial ecosystem of self-limiting bacteria using synthetic quorum-regulated lysis, Nature Microbiology 2, 17083.

14. Scott, S. R., and Hasty, J. (2016) Quorum sensing communication modules for microbial consortia, ACS Synthetic Biology 5, 969–977.

15. Young, J. W., Locke, J. C., Altinok, A., Rosenfeld, N., Bacarian, T., Swain, P. S., Mjolsness, E., and Elowitz, M. B. (2012) Measuring single-cell gene expression dynamics in bacteria using fluorescence time-lapse microscopy, Nature Protocols 7, 80.

